# HIV, Pathology and epigenetic age acceleration in different human tissues

**DOI:** 10.1101/2022.02.10.480013

**Authors:** Steve Horvath, David T.S. Lin, Michael S. Kobor, Jonathan W. Said, Susan Morgello, Elyse Singer, William H. Yong, Beth D. Jamieson, Andrew J. Levine

**Affiliations:** Department of Human Genetics, David Geffen School of Medicine, University of California Los Angeles, Los Angeles, CA 90095, USA; Department of Biostatistics, Fielding School of Public Health, University of California Los Angeles, Los Angeles, CA 90095, USA; Centre for Molecular Medicine and Therapeutics, BC Children’s Hospital Research Institute, Vancouver, Canada; Department of Pathology and Jonsson Comprehensive Cancer Center, David Geffen School of Medicine; Department of Neurology, Icahn School of Medicine at Mount Sinai, New York, NY, USA; Departments of Neuroscience and Pathology, Icahn School of Medicine at Mount Sinai, New York, NY, USA; Department of Neurology, David Geffen School of Medicine at the University of California, Los Angeles, USA; Department of Medicine, David Geffen School of Medicine at the University of California, Los Angeles, USA

## Abstract

**Background:** Epigenetic clocks based on patterns of DNA methylation have great importance in understanding aging and disease; however, there are basic questions to be resolved in their application. It remains unknown whether epigenetic age acceleration (EAA) within an individual shows strong correlation between different primary tissue sites, the extent to which tissue pathology and clinical illness correlates with EAA in the target organ, and if EAA variability across tissues differs according to sex. Considering the outsized role of age-related illness in Human Immunodeficiency Virus-1 (HIV), these questions were pursued in a sample enriched for tissue from HIV-infected individuals.

**Methods:** We used a custom methylation array to generate DNA methylation data from 661 samples representing 11 human tissues (adipose, blood, bone marrow, heart, kidney, liver, lung, lymph node, muscle, spleen, and pituitary gland) from 133 clinically-characterized, deceased individuals, including 75 infected with HIV. We developed a multimorbidity index based on the clinical disease history.

**Results:** Epigenetic age was moderately correlated across tissues. Blood had the greatest number and degree of correlation, most notably with spleen and bone marrow. However, blood did not correlate with epigenetic age of liver. EAA in liver was weakly correlated with EAA in kidney, adipose, lung, and bone marrow. Clinically, hypertension was associated with EAA in several tissues, consistent with the multi-organ impacts of this illness. HIV infection was associated with positive age acceleration in kidney and spleen. Male sex was associated with increased epigenetic acceleration in several tissues. Preliminary evidence indicates that Amyotrophic Lateral Sclerosis is associated with positive EAA in muscle tissue. Finally, greater multimorbidity was associated with greater EAA across all tissues.

**Conclusion:** Blood alone will often fail to detect EAA in other tissues. While hypertension is associated with increased EAA in several tissues, many pathologies are associated with organ-specific age acceleration.

## INTRODUCTION

Machine learning-based analyses of DNA methylation changes at cytosine residues of cytosine-phosphate-guanine dinucleotides (CpGs) have generated multivariate age predictors, known as epigenetic clocks that use specific CpG methylation levels to estimate chronological age (i.e., DNAm age) ^1-5^ and/or mortality risk ^6-8^. When studying the relationship between age-related conditions and DNAm Age, it is important to adjust the analysis for chronological age. To arrive at a non-confounded analysis, one can employ age-adjusted measures of DNAm Age, referred to as measures of epigenetic age acceleration (EAA).

The biological relevance of epigenetic measures of age acceleration can be appreciated by the fact that they relate to a host of age-related conditions and diseases ^7^. EAA has been linked to conditions such as neuropathology in the elderly ^9,10^, Down syndrome ^11^, Parkinson’s disease ^12^, Werner syndrome ^13^, physical/cognitive fitness^9^, frailty ^14^ and centenarian status ^15^. In addition to being predictive of all-cause mortality, DNAmAge acceleration in blood is associated with the risk of developing certain types of cancer ^16-19^. In older individuals, positive EAA in blood is associated with an increased risk of death from all natural causes even after accounting for known risk factors ^20-24^.

The pan tissue clock^3^ developed by the lead author in 2013 is particularly attractive for studying epigenetic aging effects in several different tissues. For example, Down syndrome is associated with strong EAA in both blood and brain tissue ^11^. Similarly, we have shown that HIV infection is associated with EAA in both blood and brain tissue^25,26^, and that age acceleration in brain is associated with HIV-associated neurocognitive disorder^10^. These results lead to natural questions: 1) does EAA in one tissue (e.g. blood) correlate with EAA in another tissue (e.g. brain), 2) is EAA in an organ associated with tissue pathology and clinical illness, 3) does EAA in specific organs underlie the higher rate of age-related illness among HIV-infected individuals, and, 4) as our previous studies have shown that males age faster than females according to an epigenetic clock analysis of blood and brain tissue, are there sex differences in EAA across tissues?

By generating a large methylation data set from 661 postmortem samples derived from 11 types of tissue, this study addresses several outstanding questions surrounding epigenetic clocks. First, to quantify the extent to which EAA in one tissue correlates with EAA in another tissue. Second, to relate measures of tissue pathology and clinical illness to EAA in the same tissue or organ. Third, to define a multimorbidity index that correlates with EAA in several non-blood tissues. Fourth, to extend previous findings from our group that HIV accelerates aging in blood and brain, by investigating the effect of HIV infection on age acceleration across these tissues. Finally, to study the effect of sex on EAA in different tissues.

## RESULTS

The mammalian methylation array platform (HorvathMammalMethylChip40) ^27^ array allowed us to calculate two different epigenetic clocks: the pan tissue clock^3^ and the skin & blood clock^5^. Since we are working with many different tissues, we primarily focus on the results from the pan tissue clock herein. As expected, the DNAm age estimate of the pan tissue clock is highly correlated with chronological age across all tissues (r=0.64, Figure 1A) and within specific tissues (Figure 1B-K). The age correlations persisted even with the somewhat limited age range of our sample, which was skewed towards older individuals (median age 57, range 26 to 91).

**Figure 1:**
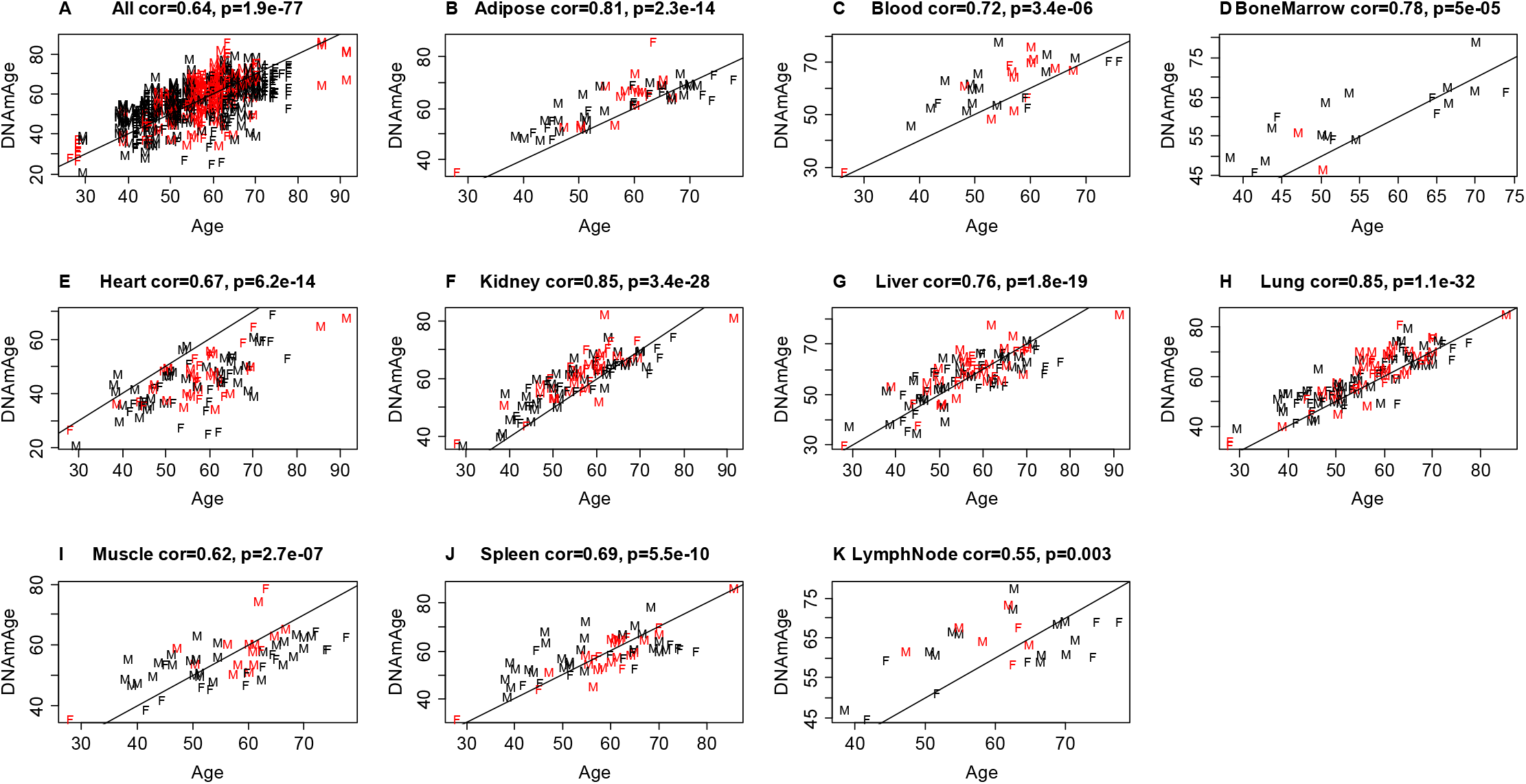
Pan tissue clock applied to different postmortem tissues. DNAmAge estimate (y-axis) versus chronological age at the time of sample collection. A) All tissues, B) adipose, C) blood, D) bone marrow E) heart, F) kidney, G) liver, H) lung, I) muscle, J) spleen, K) lymph nodes. Dots are colored by hypertension status (red=diagnosis of hypertension) and labelled by sex (F=female, M=male). The title of each plot reports the Pearson correlation coefficient. The solid line corresponds to the diagonal y=x.

### Conservation of EAA across different tissues

EAA is defined as the residual resulting from regressing DNAmAge on chronological age within the respective tissue. As illustrated in Figure 2, we found that EAA in blood is highly correlated with EAA in spleen (r=0.74), bone marrow (r=0.71), lung (r=0.62), muscle (r=0.51), adipose (r=0.47), kidney (r=0.42), and heart (r=0.34), but not in liver tissue. EAA in liver was instead correlated with EAA in kidney (r=0.49), adipose (r=0.41), lung (r=0.31) and bone marrow (r=0.30).

**Figure 2.**
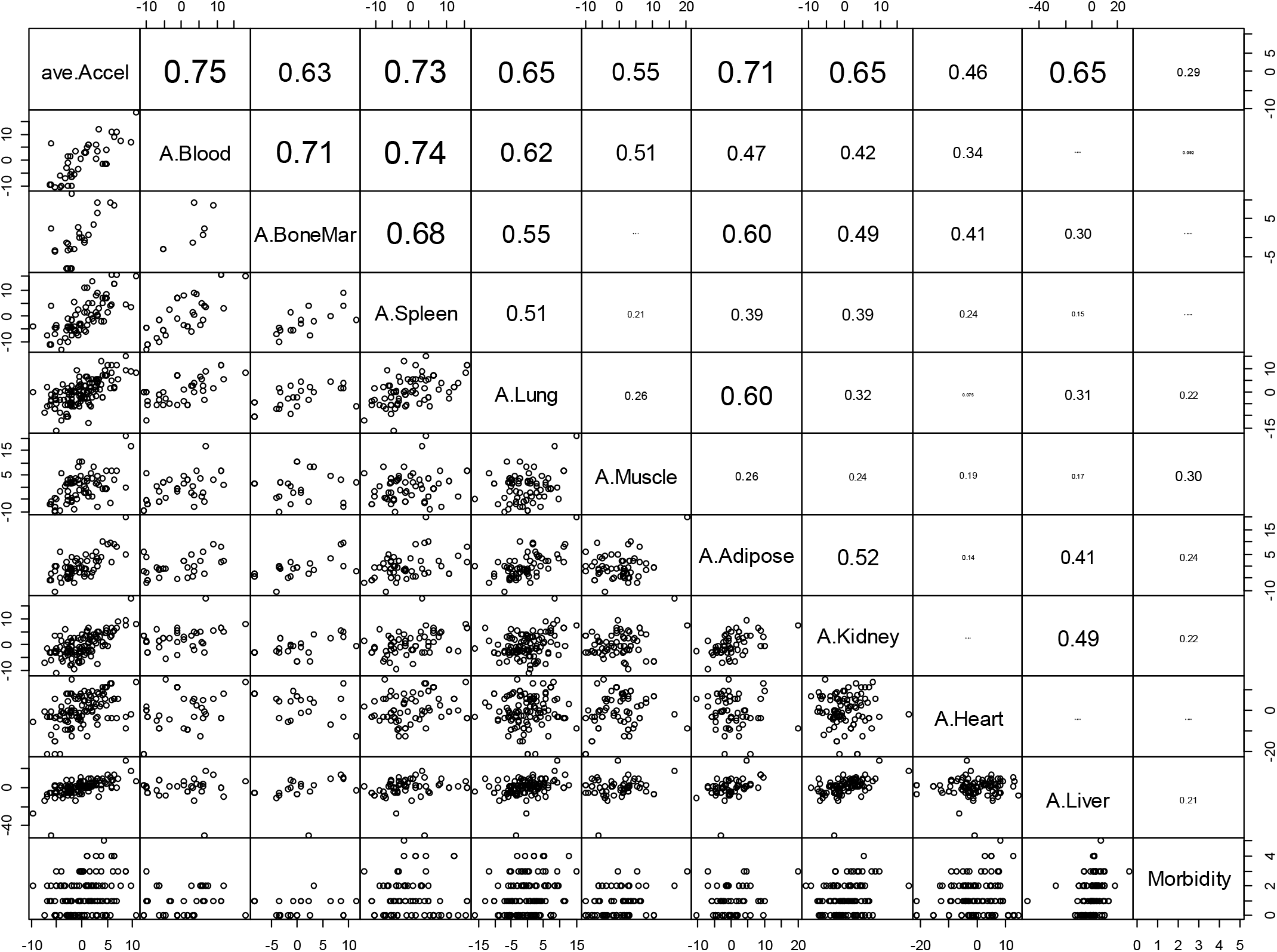
Conservation of EAA across different human tissues. The diagonal reports the respective variables for each row: EAA measures and multimorbidity index. The panels below the diagonal show the pairwise scatter plots. The numbers above the diagonal report the corresponding Pearson correlation coefficients between listed tissue for the row and the tissue listed lower down in the column. Each dot corresponds to a different person. For example, the multimorbidity index (last column) is correlated with epigenetic age acceleration in muscle (Pearson correlation 0.30). The measures of EAA were calculated within each tissue type based on the pan tissue clock. A.Blood denotes the epigenetic age acceleration in blood, i.e. the age adjusted measure of DNAmAge in blood tissue. The first variable, ave.Accel, denotes the average EAA across all tissues. Average EAA per individual, ave.Accel in the first column, was defined as average EAA across the following measures of EAA: A.Adipose, A.Blood, A.BoneMar, A.Heart, A.Kidney, A.Liver, A.Lung, A.LymphNode, A.Muscle, A.Spleen. Morbidity denotes the multimorbidity index. The analysis was limited in that each comparison involved a different set of individuals due to missing values.

### Association of EAA with and tissue/organ pathology and clinical illness

We present a detailed analysis of all measures of tissue pathology and clinical diagnoses versus EAA in lung (Supplementary Figure 7), liver (Supplementary Figure 8), heart (Supplementary Figure 9), and kidney (Supplementary Figure 10).

We also found significant associations between EAA in different tissues and hypertension, diabetes, cardiac disease, severe coronary artery disease, cerebrovascular disease, non-AIDS defining cancers, chronic renal disease, and liver disease (e.g. hepatitis or end stage liver disease) (Figure 3). Of particular interest, hypertension is associated with EAA, according to the pan tissue clock^3^, across all tissues combined (p=4.5E-5, Supplementary Figure 2A), as well as individually with kidney (p=0.0048, Supplementary Figure 2F), liver (p=0.028, Supplementary Figure 2G) and lymph nodes (p=0.033, Supplementary Figure 2K). According to the skin and blood epigenetic clock^5^, hypertension was again associated with EAA across all tissues combined (p=0.00036, Supplementary Figure 3A), and individually in kidney (p=0.0095, Supplementary Figure 3F).

**Figure 3.**
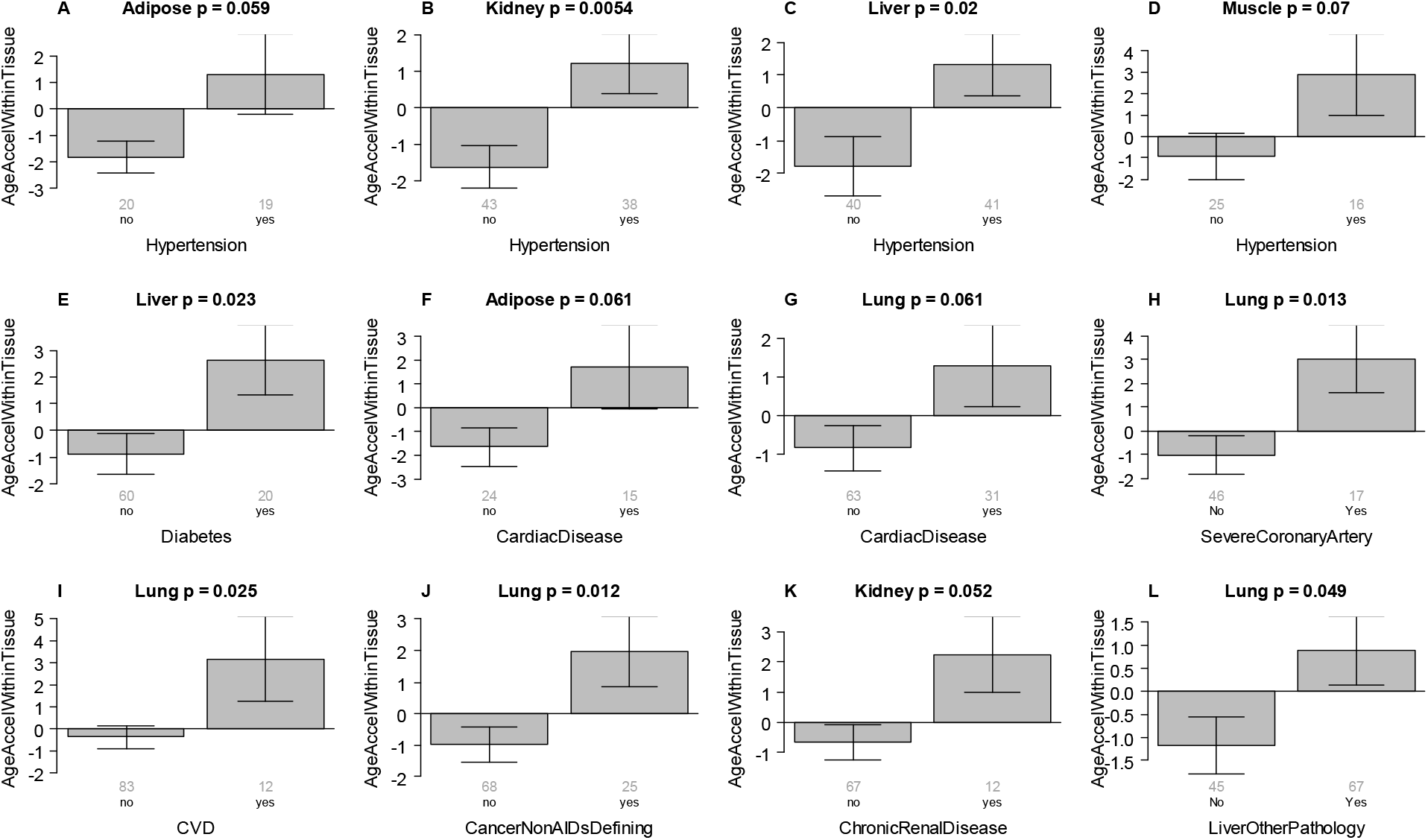
Age related conditions versus EAA in different tissues. Epigenetic age acceleration (y-axis) versus different conditions (x-axis) in different human tissues. A) Adipose, B) Kidney, C) Liver, D) Muscle, E) Liver, F) Adipose, G-J) Kung. K) Kidney. L) Lung. We caution the reader that liver-other-pathology (panel L) is not a clinical disease and may not be age related. A-D) relates hypertension status to epigenetic age acceleration. I) CVD denotes cerebrovascular disease. J) Non AIDS defining cancer status.

### Multimorbidity

For our study, we defined a novel multimorbidity index as the number of the following conditions per person: hypertension, type II diabetes, cardiovascular disease, non-AIDS defining cancer, chronic renal disease, hepatitis. For example, an individual with three conditions (e.g. hypertension, diabetes, and cardiovascular disease) was assigned a multimorbidity index score of 3. Descriptive statistics surrounding the multimorbidity index, age, HIV status, and sex are presented in Supplementary Figure 5. By design, the multimorbidity index is positively correlated with EAA in different tissues (Supplementary Figure 6). Nominally significant positive correlations (p<0.05) were observed in kidney, liver, lung, muscle (Supplementary Figure 6D-G).

Multivariable regression models that included age and sex revealed that the multimorbidity index is significantly associated with average EAA across all tissues (p=5.26E-4, Table 2). However, the average EAA, age, and sex explain only 17% of the variance in the multimorbidity index (17%, R^2=0.17). We also evaluate the relationship between multimorbidity and DNAmAge in specific tissues (Table 2). Strikingly, the pan tissue clock showed a nominally significant association (two sided p<0.062) with the multi morbidity index after correcting for age and sex.

The multimorbidity index correlated with average EAA across all tissues (r=0.29, Figure 2) and EAA in muscle (r=0.30), adipose (r=0.24), lung (r=0.22), kidney (r=0.22), liver (r=0.21) but not blood. We caution the reader that our analysis involving the multimorbidity index is biased since the conditions underlying the index were chosen so that the resulting index exhibits a positive association with EAA. Another source of bias arises from uneven missingness patterns. Figure 2 may be biased because each pairwise comparison uses different people due to missing values. Therefore, we repeated the analysis limited to 77 individuals with at most 2 missing values across a subset of tissues. We found qualitatively the same results (Supplementary Figure 4).

### HIV status and ALS status

A scatter plot reveals that kidney tissue from HIV-positive individuals exhibit increased DNAmAge compared to that of HIV-negative individuals (Supplementary Figure 1). Multivariable regression model analysis finds that HIV status is positively associated with DNAmAge in kidney (p=1.66E-4) but negatively associated with DNAmAge in muscle (p=1.16E-4) even after adjusting for the morbidity index and sex (Table 3).

Since the negative association between HIV and DNAmAge in muscle was unexpected, we carried out two follow up analyses. First, we investigated whether testosterone treatment taken by HIV-positive individuals could explain this unexpected negative association. We did not find a significant association between testosterone treatment and DNAmAge in muscle among the 20 HIV positive men for whom testosterone treatment status was available at the time of death (9 treated, 11 untreated). Second, we studied whether confounding by Amyotrophic Lateral Sclerosis (ALS) could explain the unexpected negative association. Our study involved 18 ALS cases (5 HIV positive and 13 HIV negative ALS cases). ALS was associated with significantly increased DNAmAge (5.5 years, p=0.0038) in muscle tissue even after adjusting for age and sex. However, neither ALS nor HIV status was significantly associated with DNAmAge in muscle after including both covariates in a multivariate regression model along with sex. This insignificant effect of both covariates may be due to significant multi-collinearity (chi-square test p= 5.1×10^−7^) between ALS and HIV status: most ALS cases did not have HIV and vice versa. Overall, the negative association between DNAmAge and HIV status in muscle tissue is no longer significant (p>0.2) after we adjust for ALS status.

When using the skin & blood clock^5^ for the spleen samples, we found that HIV status is associated with epigenetic age acceleration (multivariate regression model Wald test p=0.02 in model that only included Age and HIV status). The latter result echoes our original finding that HIV accelerates epigenetic age of blood since epigenetic aging effects are highly correlated between spleen and blood (r=0.74, Figure 2).

### Effect of sex on EAA across tissues/organs

Men were found to exhibit higher EAA than women across all tissues when analyzed together (p=4.4E-7, Figure 4A). Statistically significant results were also observed in individual tissues (muscle, spleen, and lymph nodes; Figure 4I-K).

**Figure 4.**
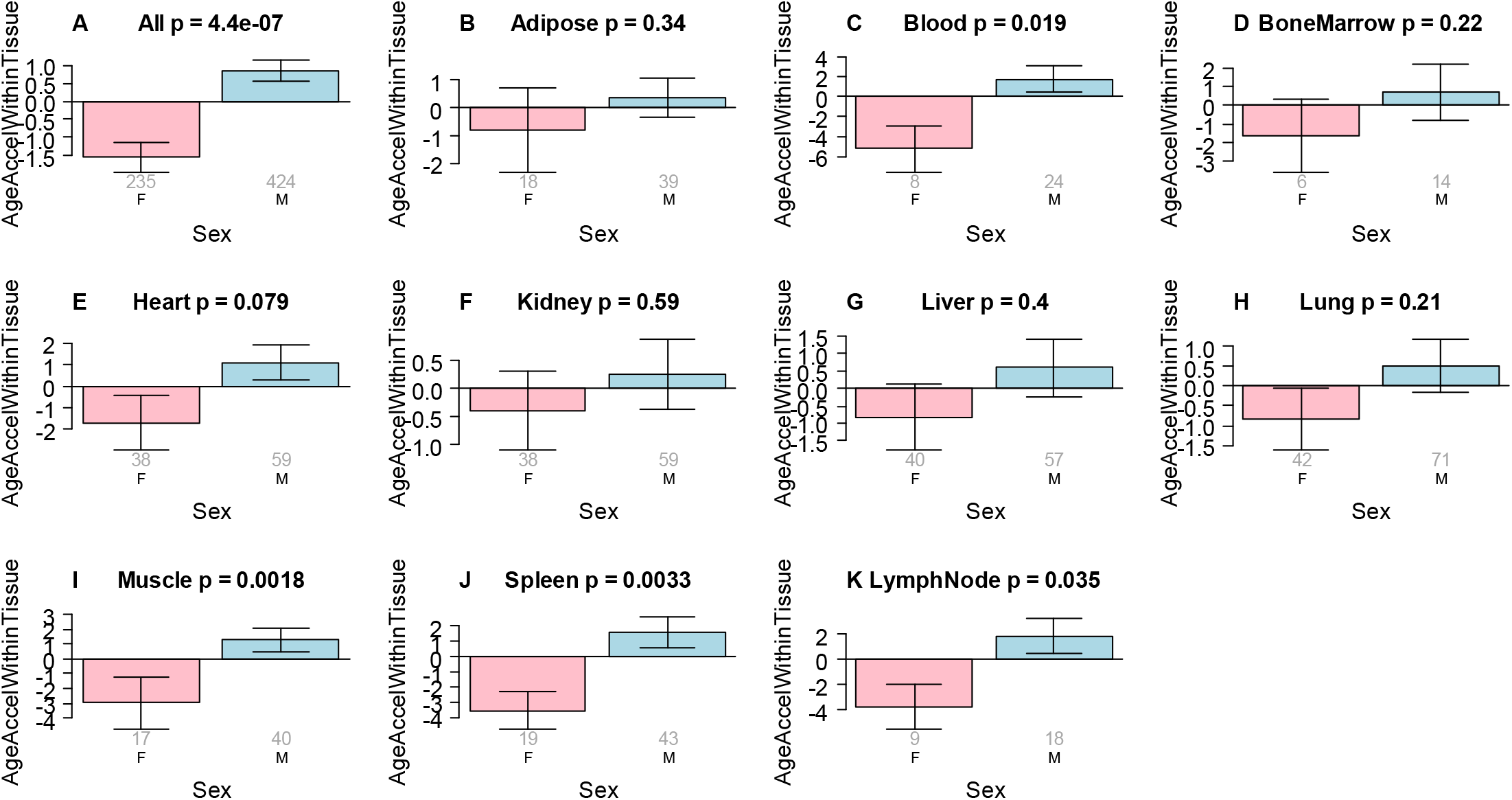
Effect of sex on EAA in different tissues. EAA within tissue (y-axis) is defined as the residual resulting from regressing DNAmAge on chronological age within a given tissue type. The grey numbers underneath each bar report the number of samples from females and males (x-axis). The title of each panel reports the result from a non-parametric group comparison test (Kruskal Wallis test). A) Results for all tissues combined, i.e. the analysis ignores tissue type. B-K report the results for different tissues/organs B) adipose, C) blood, D) bone marrow, E) heart, F) Kidney, G) liver, H) lung, I) muscle, J) spleen, K) lymph node.

## DISCUSSION

In a previous study, we showed that sex was associated with EAA in blood and brain tissue. The current study extends this finding to many other tissues ^28^. We initiated this investigation with the benefit of experience gained from our work with epigenetic clocks.^3,5,6,8^ As expected, we found that epigenetic age is largely correlated across tissues and organs. Blood had the greatest number and degree of correlations, most notably with spleen and bone marrow. Epigenetic age in blood did not correlate with that of liver. Instead, EAA in liver was weakly correlated with EAA in kidney, adipose, lung, and bone marrow. In general, heart, liver, and muscle had the weakest and fewest correlations with other tissues and organs. These findings indicate that 1) blood remains the best candidate for measuring overall EAA, and 2) the epigenetic age of tissues and organs accelerates at different rates, with some more independent than others.

In a previous study, we only found weak (but significant) positive correlations (correlation between 0.04 and 0.07) between systolic blood pressure and EAA in blood ^29^. Therefore, we were surprised to learn that hypertension was associated with EAA in several organs and tissues (heart, kidney, liver, muscle) according to the pan tissue clock. This could simply reflect the fact that hypertension leads to pathology in these organs^30^. Notably, another group recently reported that accelerated aging according to alternative epigenetic clocks is associated with organ damage in kidney, heart, brain, and peripheral arteries^31^. However, these effects were attenuated when clinical factors (BMI, diabetes, and smoking history) were included in the analysis. Conversely, inclusion of hypertension in the analyses generally did not attenuate the significance of the associations between epigenetic age and tissue pathology. Similar to our previous findings, the other research group did not find their epigenetic aging measures to be associated with diastolic or systolic blood pressure.

By design, our multimorbidity index, which is quantified as the sum of up to six medical conditions, is positively correlated with EAA in liver, lung, muscle and kidney, as well as a composite measure of age acceleration across all tissues sampled. Future studies, in which pre-mortem medical conditions are thoroughly documented, might consider applying alternative multimoribidity indexes, such as those that provide weights for more serious medical conditions^32,33^ and which have been applied in specific contexts, such as HIV research.^34-36^

Compared to tissues from uninfected individuals, those from HIV-positive individuals exhibit increased DNAmAge in kidney. HIV-1 infection is associated with an increased risk for a number of diseases and medical conditions typically associated with aging, including cardiovascular disease, osteoporosis, several cancers, kidney disease, liver disease, and cognitive decline^37-49^. We previously demonstrated the clinical relevance of DNAm-based accelerated aging in HIV-infected individuals. Specifically, we reported that the brains of deceased adults diagnosed with neurocognitive impairment within a year of death had greater age acceleration than those who were cognitively normal^50^. More recently, we reported that neurodevelopment and neuropsychological deficits in perinatally-HIV-infected adolescents are associated with accelerated aging^26,51^. The current findings support age acceleration only in kidney, as well as an association between age acceleration and kidney disease. However, the lack of findings of relatively greater age acceleration in other tissues does not support the hypothesis that HIV-induced accelerated aging leads to increased incidence of age-related medical conditions via direct effects on underlying organs.

The unexpected finding that HIV status is negatively associated with DNAmAge in muscle (even after adjusting for the morbidity index and sex) probably reflects confounding by ALS status. In the same data, we found that ALS was associated with significant positive age acceleration in muscle tissue. Strictly speaking, strong confounding/multi-collinearity between ALS status and HIV status does not allow us to distinguish between the following possibilities: either ALS is associated with positive age acceleration or HIV is associated with negative age acceleration in muscle. Future studies focusing on these aspects will be needed to resolve the observed complexity.

While this study reveals conservation of epigenetic aging effects across different tissues, we expect that blood will only be a suboptimal surrogate for other tissues when it comes to epigenetic aging effects. This judgement is based on a) the moderate correlation coefficients between age acceleration in blood and that of other tissues and b) the fact that several conditions are associated with tissue specific age acceleration effects. It will be advisable to profile several sources of DNA (including blood, buccal cells, adipose, skin) to get a comprehensive picture of the epigenetic aging state of an individual.

## METHODS

### Tissue Samples

This study was conducted in accordance with the University of California, Los Angeles Medical Institutional Review Board (IRB). Clinical and pathological data and biological samples came from 133 HIV-infected and uninfected individuals enrolled in either the National Neurological AIDS Bank (NNAB) or Manhattan HIV Brain Bank MHBB) sites of the National NeuroAIDS Tissue Consortium (NNTC)^52,53^. All individuals died between 2001 and 2016. These biorepositories operate in accordance with their local IRBs and act as “honest brokers” in maintaining participant confidentiality. All samples were obtained with approved consents allowing for genetic analysis. Donors contributing to this study died between 2001 and 2016.

### Tissue Pathology

At the time of autopsy, representative samples of all tissues were obtained and flash frozen, as well as fixed in formalin. Formalin fixed tissues were processed for paraffin embedding and routine histology. Hematoxylin and eosin stains were examined by board certified anatomic pathologists (SM & JS), and special stains obtained as indicated by the histopathology. Slides were reviewed by two pathologists to arrive at concordant diagnoses.

### Clinical Characterization

Age and sex of donors for each tissue are displayed in Table 1. Medical diagnoses were obtained by self-report and/or medical record review. The NNTC routinely collects data on hypertension, diabetes, dyslipidemia, hepatitis and liver disease, chronic renal disease, cardiac disease, chronic obstructive pulmonary disease, cancer, cerebrovascular disease, and diverse neurologic conditions.

**Table 1.** Age and sex of persons from which tissue samples were derived

| Tissue | Total N | Female N | Mean Age | Min. Age | Max. Age |
| --- | --- | --- | --- | --- | --- |
| Adipose | 57 | 18 | 57.3 | 27.9 | 77.6 |
| Blood | 32 | 8 | 55 | 26.2 | 76.1 |
| Bone Marrow | 20 | 6 | 54.7 | 38.3 | 73.9 |
| Heart | 97 | 38 | 56.1 | 27.9 | 91.2 |
| Kidney | 97 | 38 | 55.7 | 27.9 | 91.2 |
| Liver | 97 | 40 | 56.2 | 27.9 | 91.2 |
| Lung | 113 | 42 | 55.7 | 27.9 | 85.3 |
| Lymph Node | 27 | 9 | 60.2 | 38.3 | 77.6 |
| Muscle | 57 | 17 | 56.7 | 27.9 | 77.6 |
| Pituitary | 2 | 0 | 54.3 | 43.7 | 64.9 |
| Spleen | 62 | 19 | 57.3 | 27.9 | 85.3 |

**Table 2.**
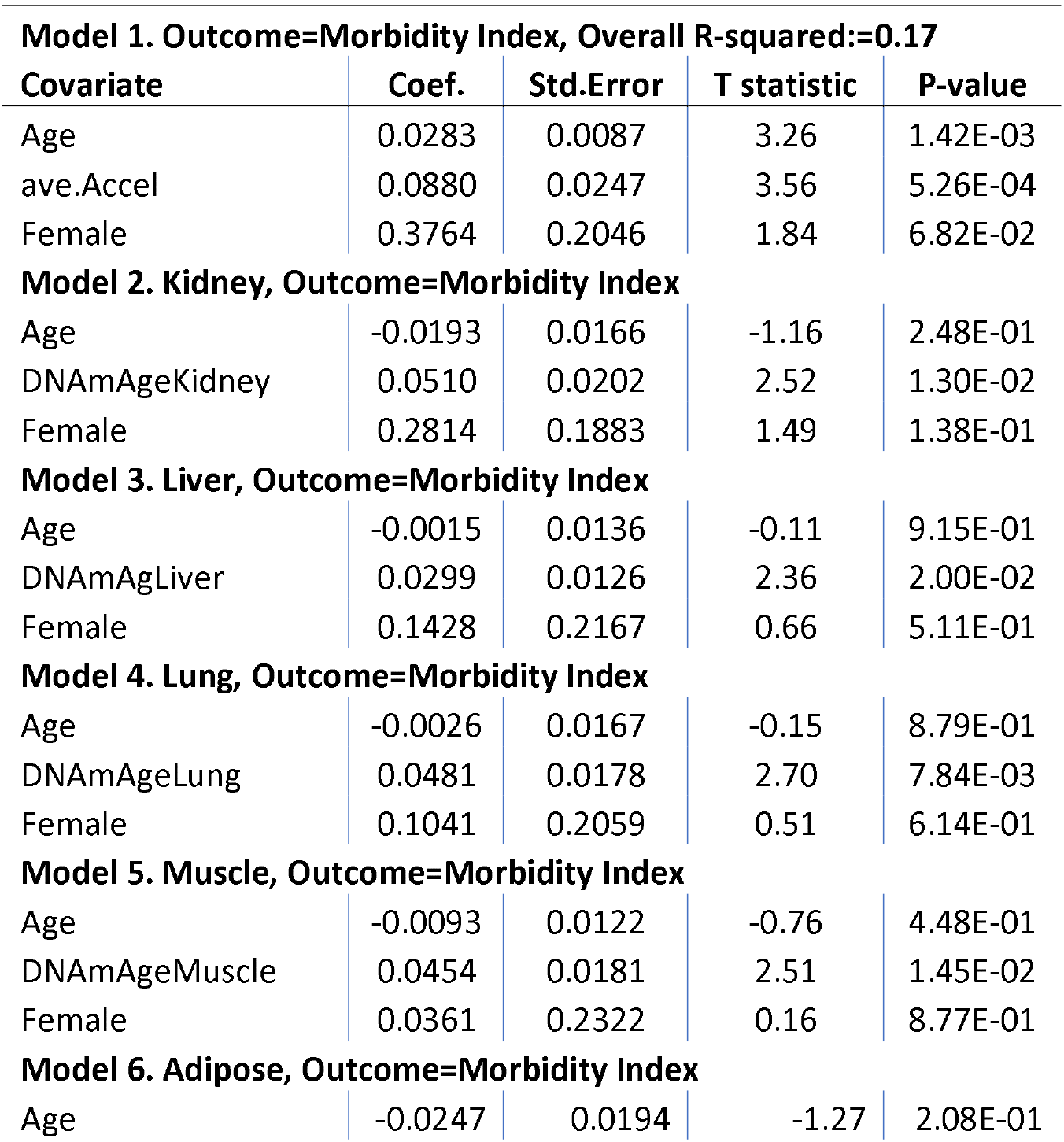
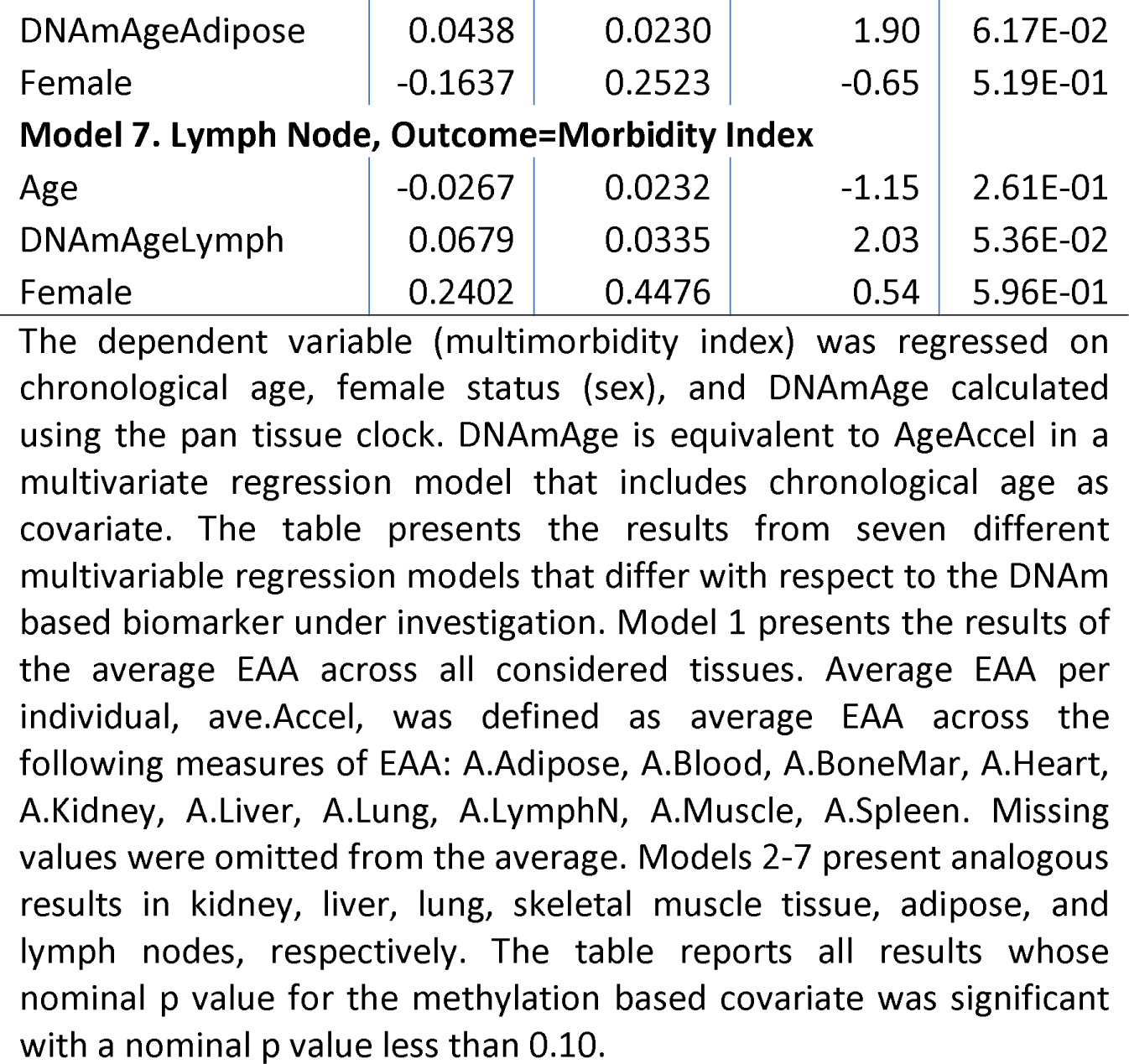
Multivariable regression model of the multimorbidity index

**Model 1. Outcome=Morbidity Index, Overall R-squared:=0.17**
| Covariate | Coef. | Std.Error | T statistic | P-value |
| --- | --- | --- | --- | --- |
| Age | 0.0283 | 0.0087 | 3.26 | 1.42E-03 |
| ave.Accel | 0.0880 | 0.0247 | 3.56 | 5.26E-04 |
| Female | 0.3764 | 0.2046 | 1.84 | 6.82E-02 |
| **Model 2. Kidney, Outcome=Morbidity Index** |  |  |  |  |
| Age | -0.0193 | 0.0166 | -1.16 | 2.48E-01 |
| DNAMAgeKidney | 0.0510 | 0.0202 | 2.52 | 1.30E-02 |
| Female | 0.2814 | 0.1883 | 1.49 | 1.38E-01 |
| **Model 3. Liver, Outcome=Morbidity Index** |  |  |  |  |
| Age | -0.0015 | 0.0136 | -0.11 | 9.15E-01 |
| DNAMAgLiver | 0.0299 | 0.0126 | 2.36 | 2.00E-02 |
| Female | 0.1428 | 0.2167 | 0.66 | 5.11E-01 |
| **Model 4. Lung, Outcome=Morbidity Index** |  |  |  |  |
| Age | -0.0026 | 0.0167 | -0.15 | 8.79E-01 |
| DNAMAgeLung | 0.0481 | 0.0178 | 2.70 | 7.84E-03 |
| Female | 0.1041 | 0.2059 | 0.51 | 6.14E-01 |
| **Model 5. Muscle, Outcome=Morbidity Index** |  |  |  |  |
| Age | -0.0093 | 0.0122 | -0.76 | 4.48E-01 |
| DNAMAgeMuscle | 0.0454 | 0.0181 | 2.51 | 1.45E-02 |
| Female | 0.0361 | 0.2322 | 0.16 | 8.77E-01 |
| **Model 6. Adipose, Outcome=Morbidity Index** |  |  |  |  |
| Age | -0.0247 | 0.0194 | -1.27 | 2.08E-01 |
| DNAmAgeAdipose | 0.0438 | 0.0230 | 1.90 | 6.17E-02 |
| Female | -0.1637 | 0.2523 | -0.65 | 5.19E-01 |
| **Model 7. Lymph Node, Outcome=Morbidity Index** |  |  |  |  |
| Age | -0.0267 | 0.0232 | -1.15 | 2.61E-01 |
| DNAmAgeLymph | 0.0679 | 0.0335 | 2.03 | 5.36E-02 |
| Female | 0.2402 | 0.4476 | 0.54 | 5.96E-01 |

The dependent variable (multimorbidity index) was regressed on chronological age, female status (sex), and DNAmAge calculated using the pan tissue clock. DNAmAge is equivalent to AgeAccel in a multivariate regression model that includes chronological age as covariate. The table presents the results from seven different multivariable regression models that differ with respect to the DNAm based biomarker under investigation. Model 1 presents the results of the average EAA across all considered tissues. Average EAA per individual, ave.Accel, was defined as average EAA across the following measures of EAA: A.Adipose, A.Blood, A.BoneMar, A.Heart, A.Kidney, A.Liver, A.Lung, A.LymphN, A.Muscle, A.Spleen. Missing values were omitted from the average. Models 2-7 present analogous results in kidney, liver, lung, skeletal muscle tissue, adipose, and lymph nodes, respectively. The table reports all results whose nominal p value for the methylation based covariate was significant with a nominal p value less than 0.10.

**Table 3.**
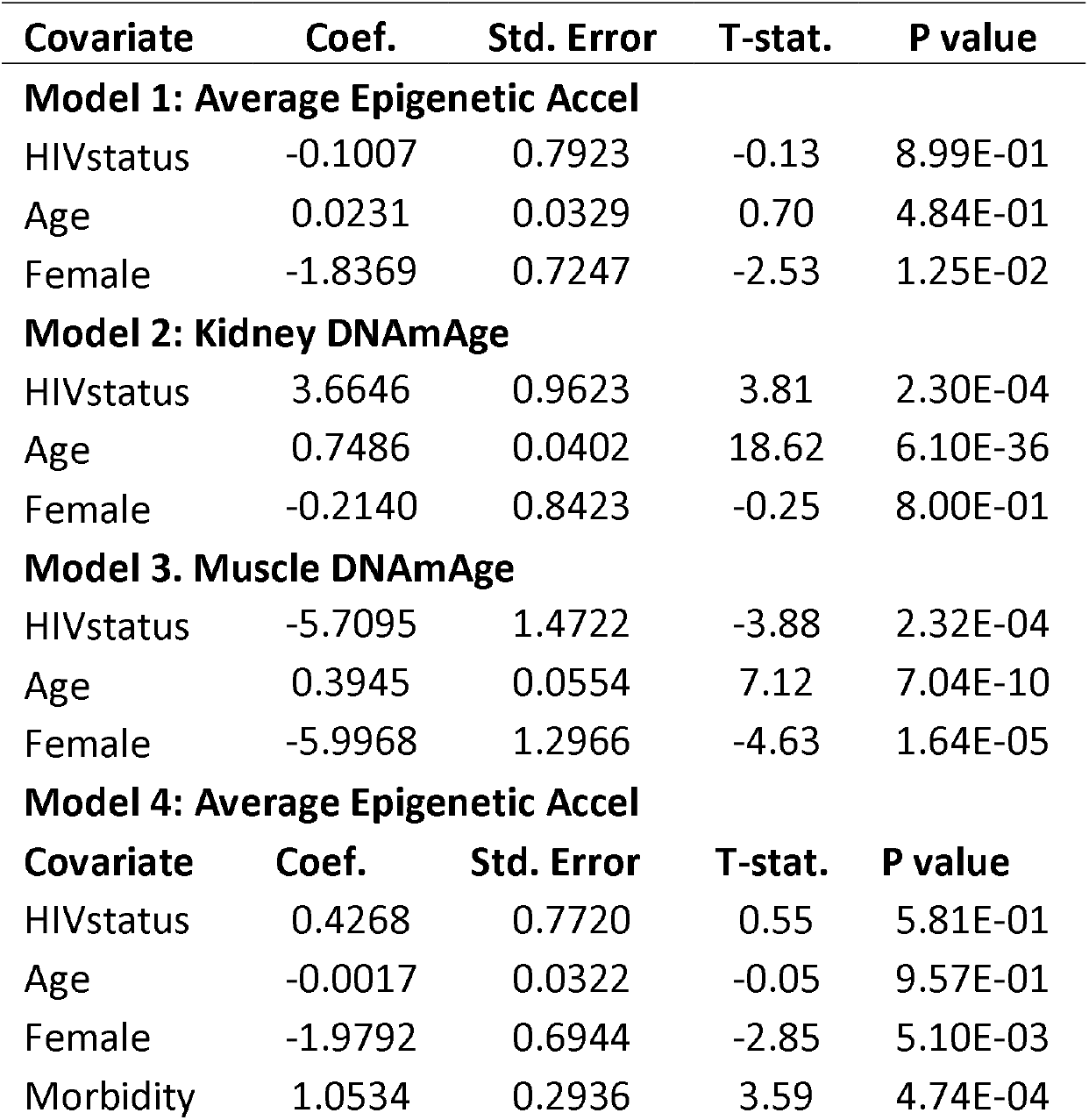
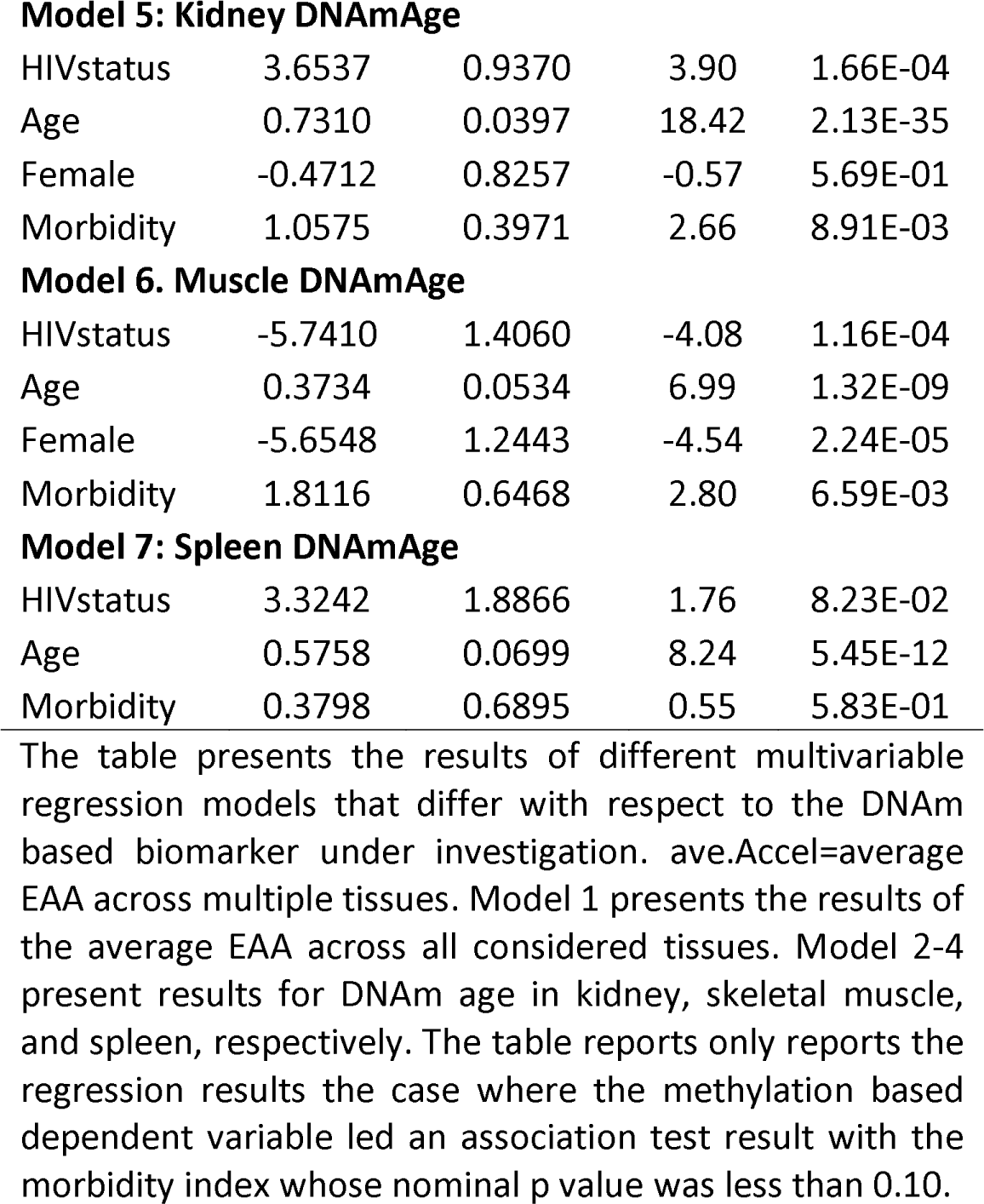
Multivariable regression model of HIV status

| Covariate | Coef. | Std. Error | T-stat. | P value |
| --- | --- | --- | --- | --- |
| **Model 1: Average Epigenetic Accel** |  |  |  |  |
| HIVstatus | -0.1007 | 0.7923 | -0.13 | 8.99E-01 |
| Age | 0.0231 | 0.0329 | 0.70 | 4.84E-01 |
| Female | -1.8369 | 0.7247 | -2.53 | 1.25E-02 |
| **Model 2: Kidney DNAmAge** |  |  |  |  |
| HIVstatus | 3.6646 | 0.9623 | 3.81 | 2.30E-04 |
| Age | 0.7486 | 0.0402 | 18.62 | 6.10E-36 |
| Female | -0.2140 | 0.8423 | -0.25 | 8.00E-01 |
| **Model 3: Muscle DNAmAge** |  |  |  |  |
| HIVstatus | -5.7095 | 1.4722 | -3.88 | 2.32E-04 |
| Age | 0.3945 | 0.0554 | 7.12 | 7.04E-10 |
| Female | -5.9968 | 1.2966 | -4.63 | 1.64E-05 |
| **Model 4: Average Epigenetic Accel** |  |  |  |  |
| Covariate | Coef. | Std. Error | T-stat. | P value |
| HIVstatus | 0.4268 | 0.7720 | 0.55 | 5.81E-01 |
| Age | -0.0017 | 0.0322 | -0.05 | 9.57E-01 |
| Female | -1.9792 | 0.6944 | -2.85 | 5.10E-03 |
| Morbidity | 1.0534 | 0.2936 | 3.59 | 4.74E-04 |
| **Model 5: Kidney DNAmAge** |  |  |  |  |
| HIVstatus | 3.6537 | 0.9370 | 3.90 | 1.66E-04 |
| Age | 0.7310 | 0.0397 | 18.42 | 2.13E-35 |
| Female | -0.4712 | 0.8257 | -0.57 | 5.69E-01 |
| Morbidity | 1.0575 | 0.3971 | 2.66 | 8.91E-03 |
| **Model 6: Muscle DNAmAge** |  |  |  |  |
| HIVstatus | -5.7410 | 1.4060 | -4.08 | 1.16E-04 |
| Age | 0.3734 | 0.0534 | 6.99 | 1.32E-09 |
| Female | -5.6548 | 1.2443 | -4.54 | 2.24E-05 |
| Morbidity | 1.8116 | 0.6468 | 2.80 | 6.59E-03 |
| **Model 7: Spleen DNAmAge** |  |  |  |  |
| HIVstatus | 3.3242 | 1.8866 | 1.76 | 8.23E-02 |
| Age | 0.5758 | 0.0699 | 8.24 | 5.45E-12 |
| Morbidity | 0.3798 | 0.6895 | 0.55 | 5.83E-01 |

The table presents the results of different multivariable regression models that differ with respect to the DNAm based biomarker under investigation. ave.Accel=average EAA across multiple tissues. Model 1 presents the results of the average EAA across all considered tissues. Model 2-4 present results for DNAm age in kidney, skeletal muscle, and spleen, respectively. The table reports only reports the regression results the case where the methylation based dependent variable led an association test result with the morbidity index whose nominal p value was less than 0.10.

### Multimorbidity

The conditions underlying the multimorbidity index were chosen so that the resulting index would exhibit a positive association between the index and EAA. In other words, the analysis is biased. Thus, the p-value should be interpreted as descriptive measure as opposed to an inferential measure. Our version of the multimorbidity index counts the number of conditions an individual had at the time of death. The selected conditions were: Hypertension, Type-2 Diabetes, Cardiovascluar disease (CVD), Non-AIDS Defining Cancers, Chronic Renal Disease, and Hepatitis.

### DNA Methylation

We used the mammalian methylation array ^27^ to generate DNA methylation data from n=672 samples representing 11 human tissues (adipose, blood, bone marrow, heart, kidney, liver, lung, lymph node, muscle, and spleen, pituitary gland) (Table 1). We removed n=11 samples from the analysis since they were outliers according to hierarchical clustering, the inter-array correlation coefficient, or detection p values.

All methylation data were generated using a custom Illumina methylation array (HorvathMammalMethylChip40) based on 37492 CpG sites ^27^. Of these, 1986 CpGs were cgiseb based on their utility for human biomarker studies; these CpGs, which were previously implemented in human Illumina Infinium arrays (EPIC, 450K, 27K), were selected due to their relevance for estimating human age, human blood cell counts or the proportion of neurons in human brain tissue. The particular subset of species for each probe is provided in the chip manifest file at the NCBI Gene Expression Omnibus (GEO) platform (GPL28271). The “noob” normalization method was used to define beta values using the *minfi* R package ^54,55^.

## ACKNOWLEDGEMENTS and FUNDING

This study was funded by the National Institute on Aging grant R21AG046954 (Horvath & Levine), NIA 1U01AG060908 (Horvath) and Paul G. Allen Frontiers Group (SH). The NNAB and MHBB are funded by the National Institute of Mental Health grants U24MH100929 (Singer) and U24MH100931 (Morgello), respectively. BDJ was also supported by R01 AG052340 NIA. These biorepositories thank their staff and study participants for their time and dedication.

## CONTRIBUTIONS

SH and AJL conceived of the study. SH carried out most of the statistical analyses. DTSL and MK processed the samples for DNA methylation. SM, JS, and WY conducted pathological characterization. All authors contributed data, helped with the write up, and participated in the interpretation of the results.

## CORRESPONDING AUTHOR

Correspondence to Steve Horvath

## Supplementary Figures

**Supplementary Figure 1.**
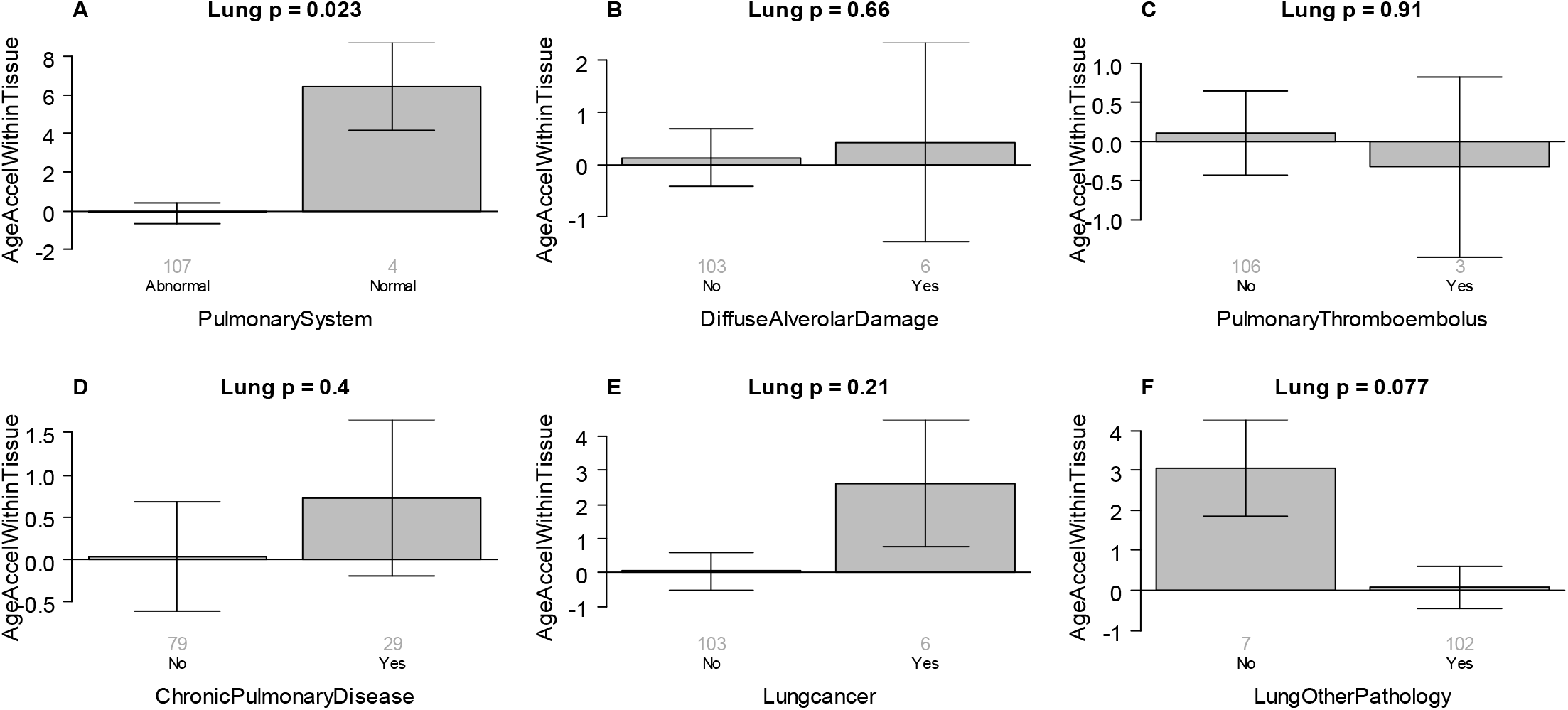
Detailed analysis of lung tissue. Epigenetic age acceleration (y-axis) versus various conditions (x-axis). Each barplot reports a two sided Kruskal Wallis test p value. Each bar depicts mean values and one standard error. Group sizes (counts) are reported under each bar. Epigenetic age acceleration (y-axis) versus A) abnormal plumonary system status, B) diffuse alverolar damaage, C) pulmonary thromboembolus, D) chronic pulmonary disease, E) lung cancer, F) other pathology.

**Supplementary Figure 2.**
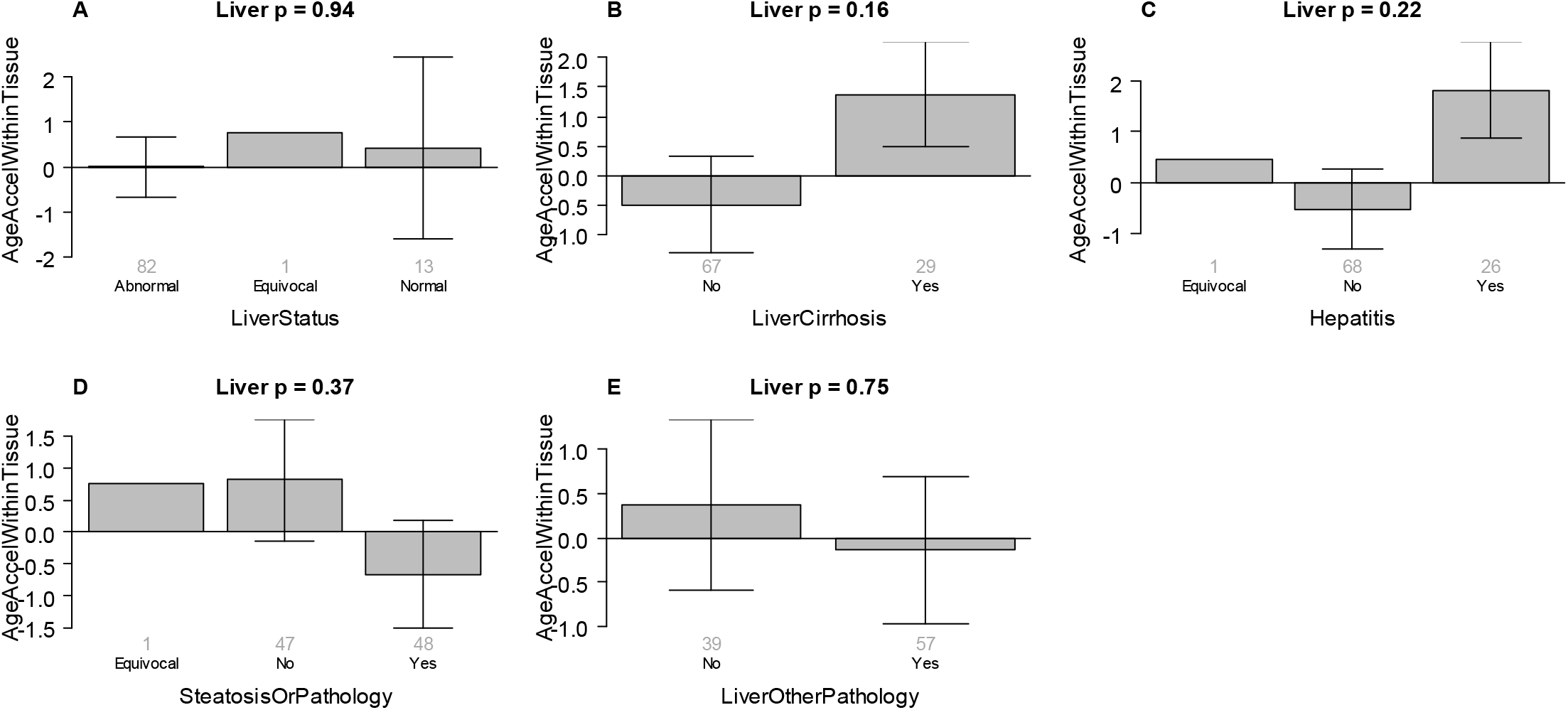
Detailed analysis of liver. Epigenetic age acceleration (y-axis) versus various conditions (x-axis). Each barplot reports a two sided Kruskal Wallis test p value. Each bar depicts mean values and one standard error. Group sizes (counts) are reported under each bar.

**Supplementary Figure 3.**
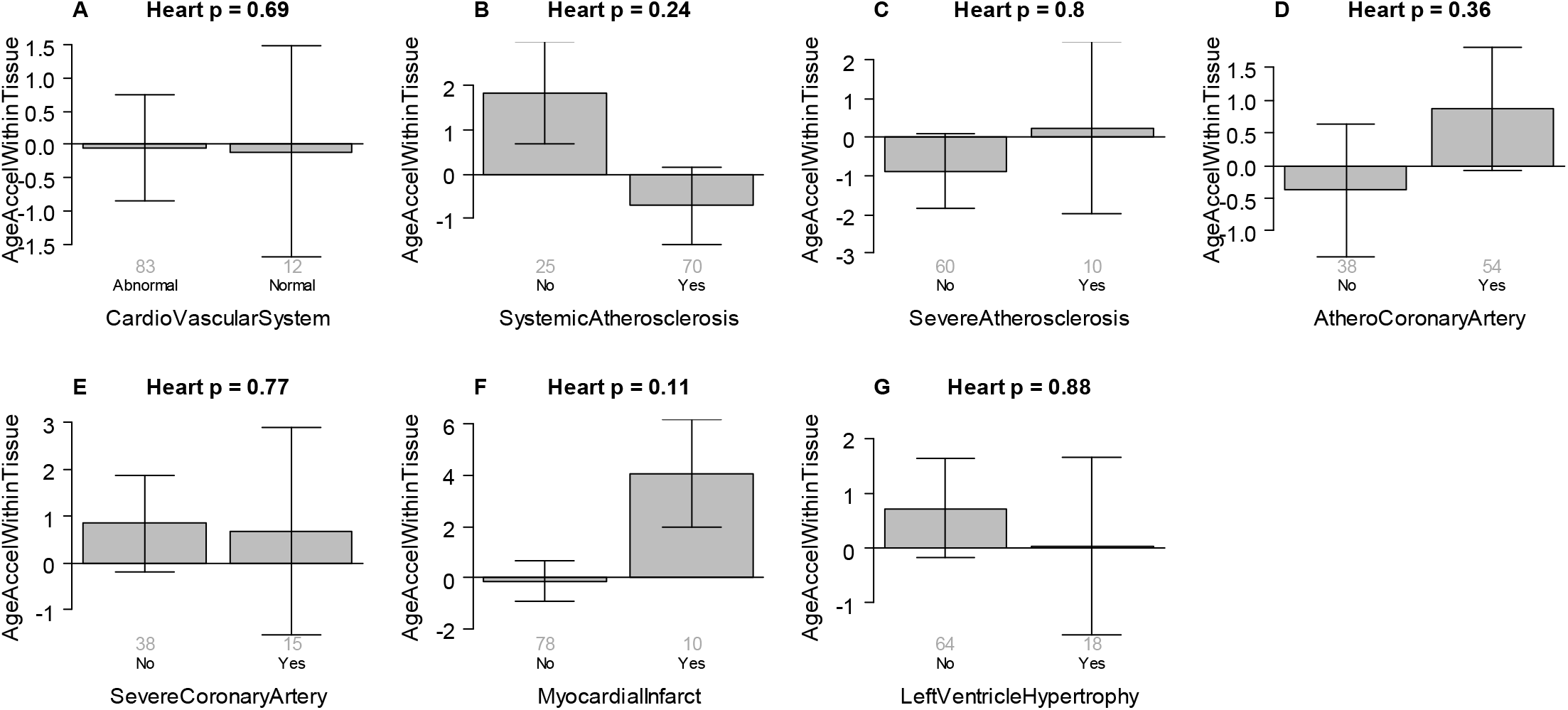
Detailed analysis of aorta and heart. Epigenetic age acceleration in the heart (y-axis) versus various conditions (x-axis). Some of the conditions pertain to the aorta (e.g. atherosclerosis in panels B,C). Panel E is a subset of those who have a Yes value in panel D. Each barplot reports a two sided Kruskal Wallis test p value. Each bar depicts mean values and one standard error. Group sizes (counts) are reported under each bar.

**Supplementary Figure 4.**
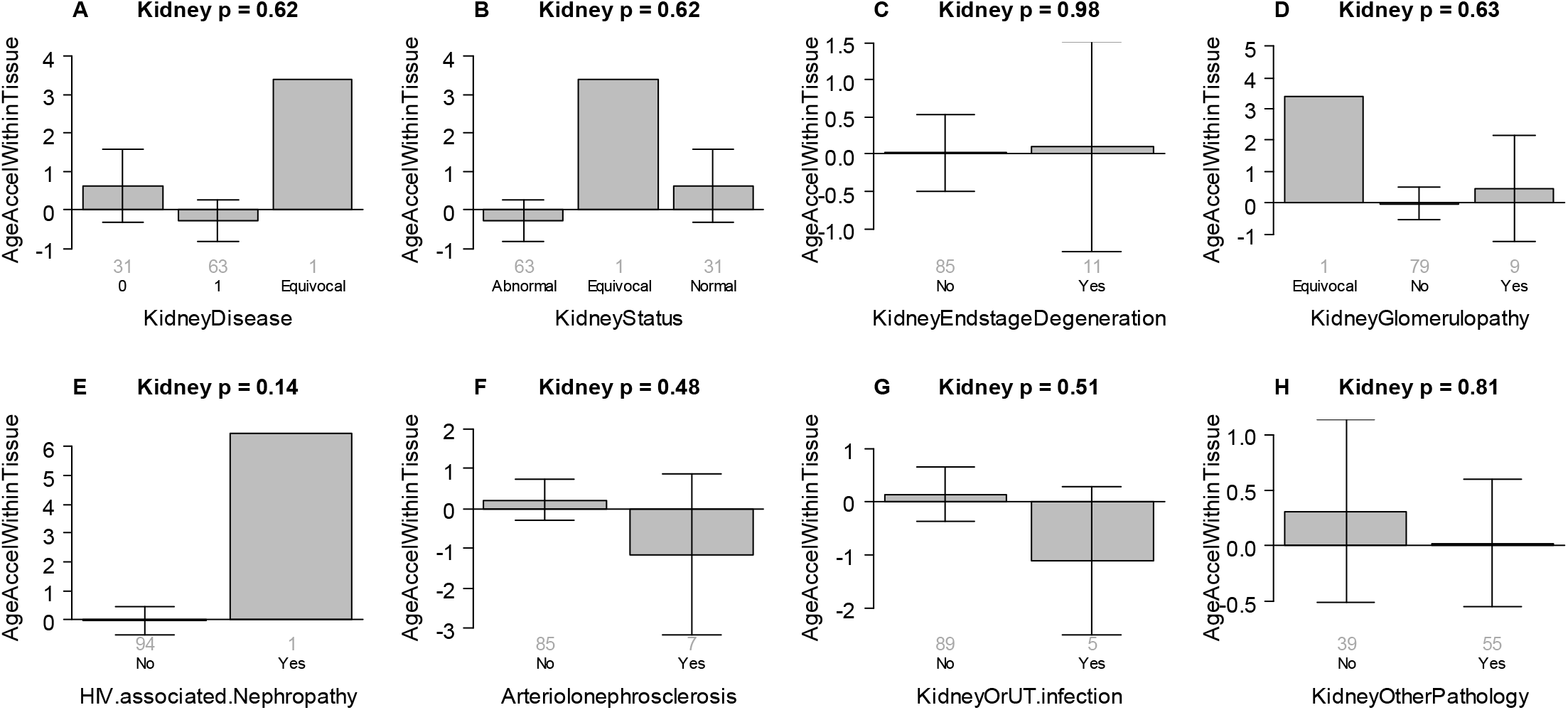
Detailed analysis of heart. A) X axis labels 0 and 1 refer to no and yes respectively. Epigenetic age acceleration in kidney (y-axis) versus various conditions (x-axis). Each barplot reports a two sided Kruskal Wallis test p value. Each bar depicts mean values and one standard error. Group sizes (counts) are reported under each bar.

**Supplementary Figure 5.**
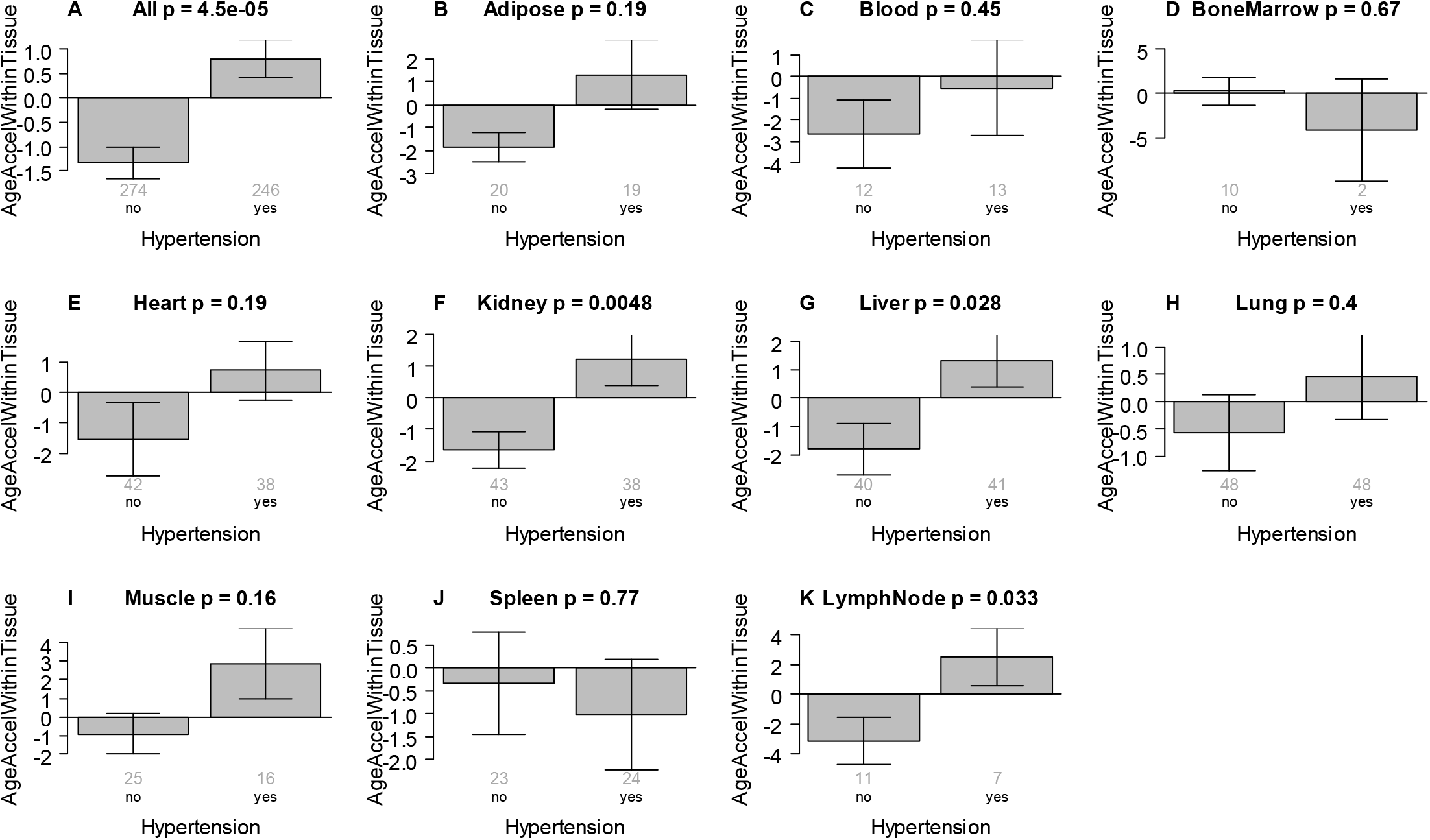
Hypertension status versus EAA of the pan tissue clock. EAA was defined as residual resulting from regressing DNAmAge on chronological age within the respective tissue. Hypertension status (x-axis) versus EAA in A) all tissues, B) adipose, C) blood, D) bone marrow E) heart, F) kidney, G) liver, H) lung, I) muscle, J) spleen, K) lymph nodes. The title of each panel reports a Kruskal Wallis test p-value. The bar plots report the mean value and one standard error. The small grey numbers under each bar report group sizes.

**Supplementary Figure 6.**
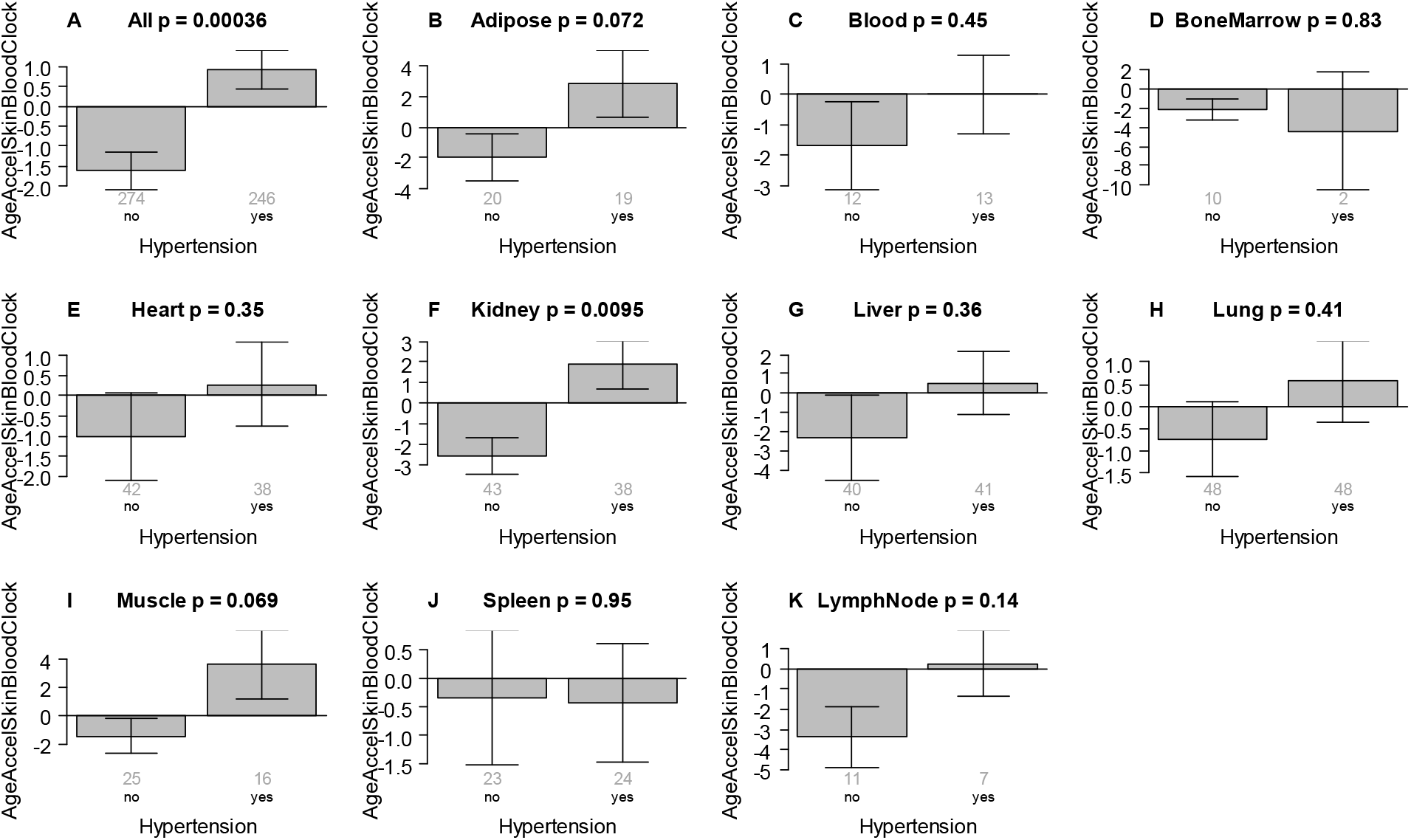
Hypertension versus EAA according to the skin and blood clock. EAA based on the pan tissue clock from Horvath 2018 (y-axis) versus hypertension status (x-axis) in A) all tissues, B) adipose, C) blood, D) bone marrow E) heart, F) kidney, G) liver, H) lung, I) muscle, J) spleen, K) lymph nodes. The title of each panel reports a Kruskal Wallis test p-value. The bar plots report the mean value and one standard error. The small grey numbers under each bar report group sizes. EAA was defined as residual resulting from regressing DNAmAge on chronological age within the respective tissue.

**Supplementary Figure 7.**
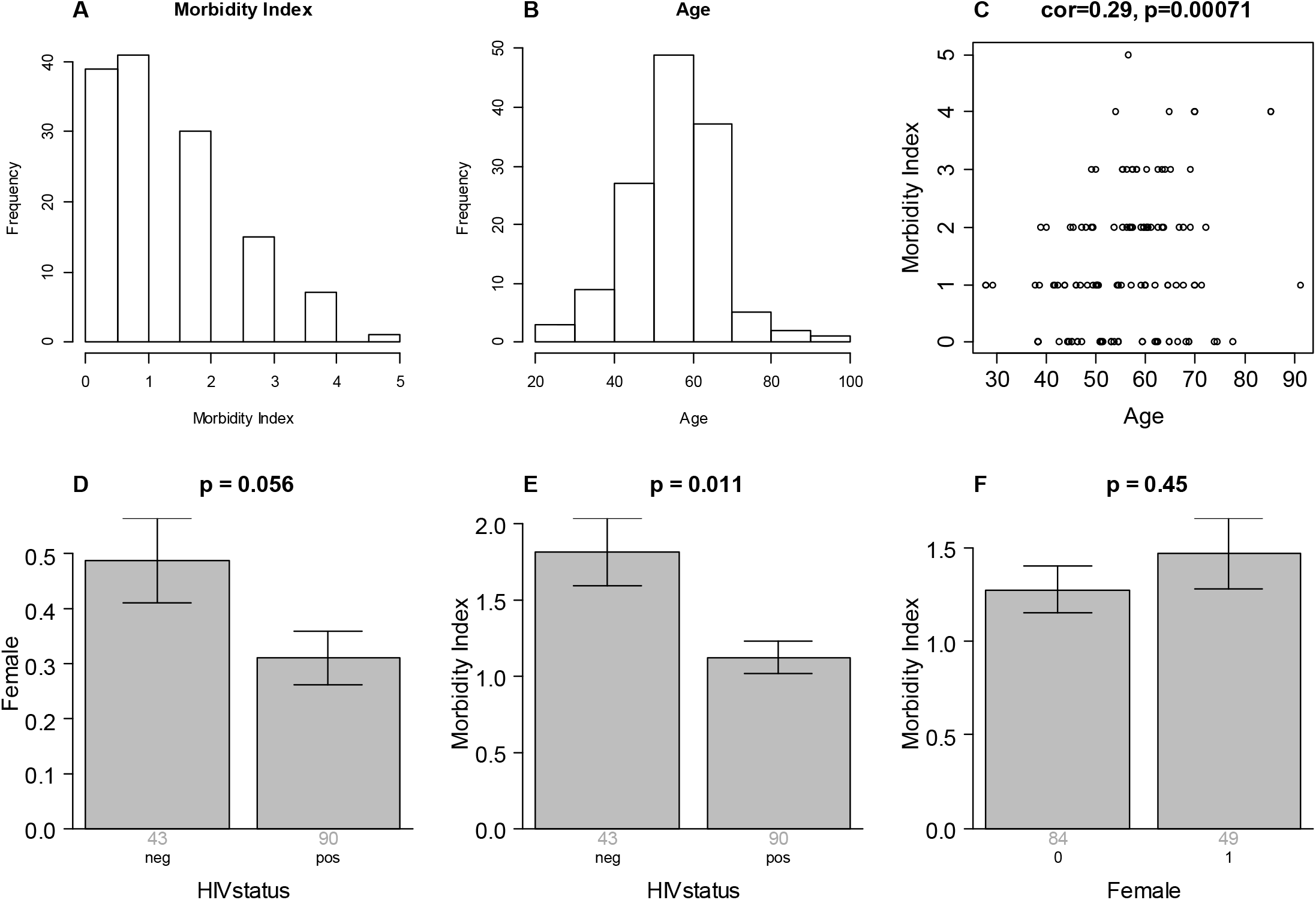
Descriptive statistics surrounding the multimorbidity index versus, age, HIV status, and sex. A) Histogram of the multimorbidity index. B) Histogram of age, C) Multimorbidity index versus age, D) Proportion of females versus HIV grouping status. Multimorbidity index versus E) HIV status, and F) female status (1=female). We caution the reader that these marginal associations are confounded. For example, the relationship between the morbidity index and HIV status is confounded by age and sex. For this reason, we use multivariable regression models.

**Supplementary Figure 8.**
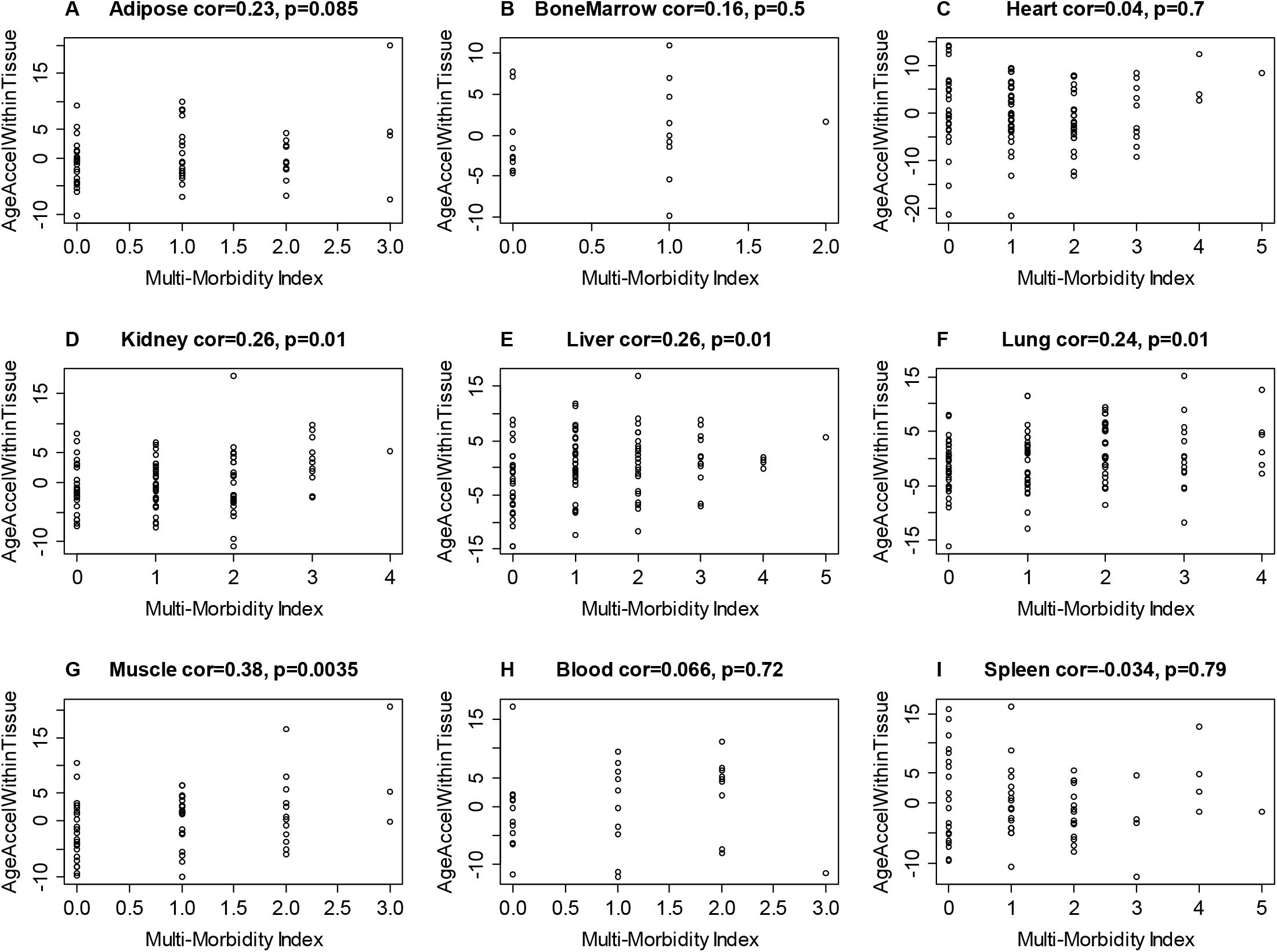
Multimorbidity index versus EAA in different tissues. Each panel reports the tissue/organ, the Pearson correlation coefficient and corresponding p value.

**Supplementary Figure 9.**
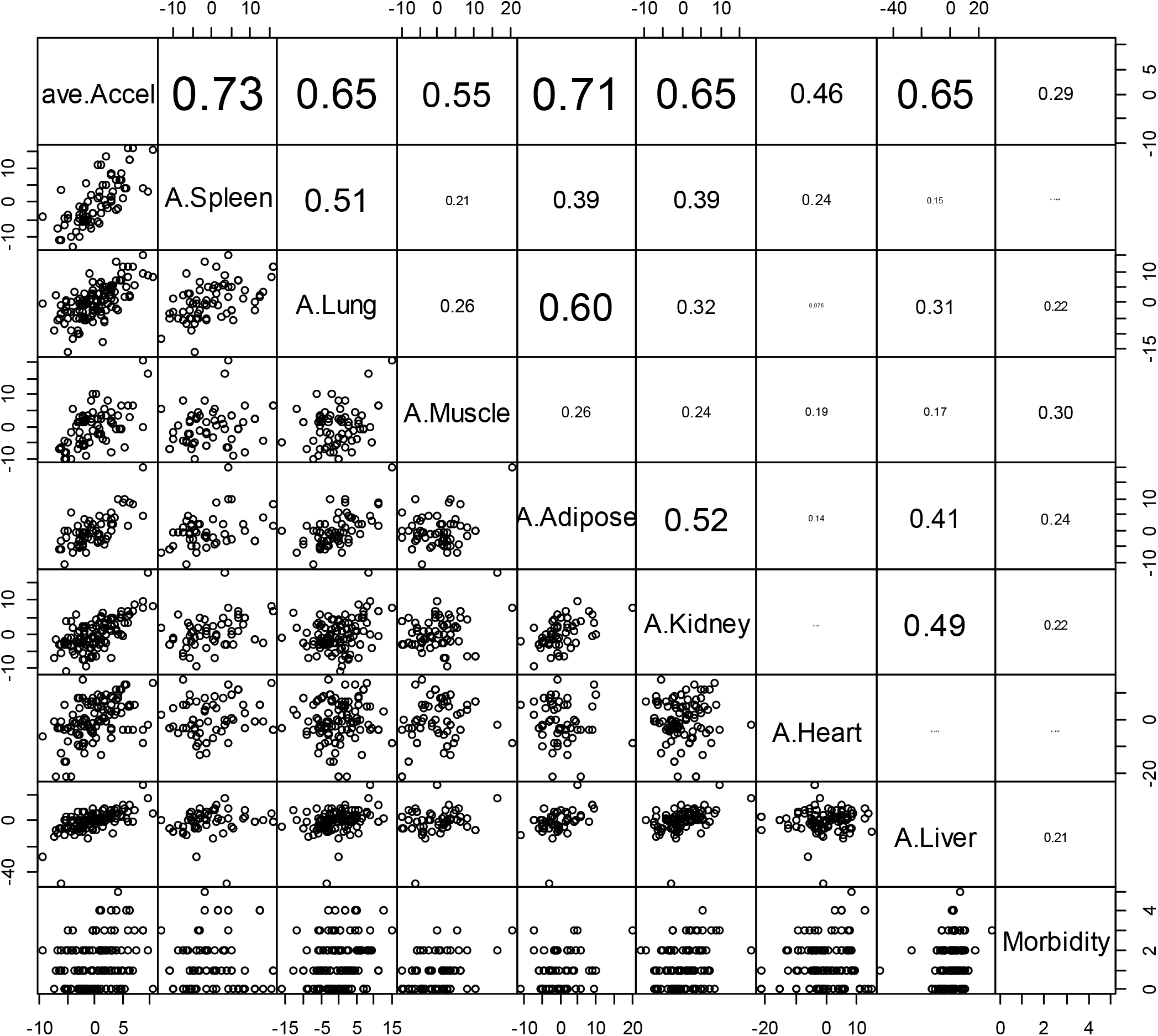
Conservation of EAA across different tissues limited to individuals with few missing values. This is the analog of Figure 3 but it involves only n=77 individuals with at most 2 missing value per measure of EAA. The diagonal reports the respective variables: EAA measures and multimorbidity index. The panels below the diagonal show the pairwise scatter plots. The numbers above the diagonal report the corresponding Pearson correlation coefficients. Each dot corresponds to a different person. The measures of EAA were calculated within each tissue type based on the pan tissue clock (Horvath 2013). The first variable, ave.Accel, denotes the average EAA across all tissues. Average EAA per individual, ave.Accel, was defined as average EAA across the following measures of EAA: A.Adipose, A.Blood, A.BoneMar, A.Heart, A.Kidney, A.Liver, A.Lung, A.LymphN, A.Muscle, A.Spleen. Here A.Blood denotes EAA in blood. Morbidity denotes the multimorbidity index.

**Supplementary Figure 10.**
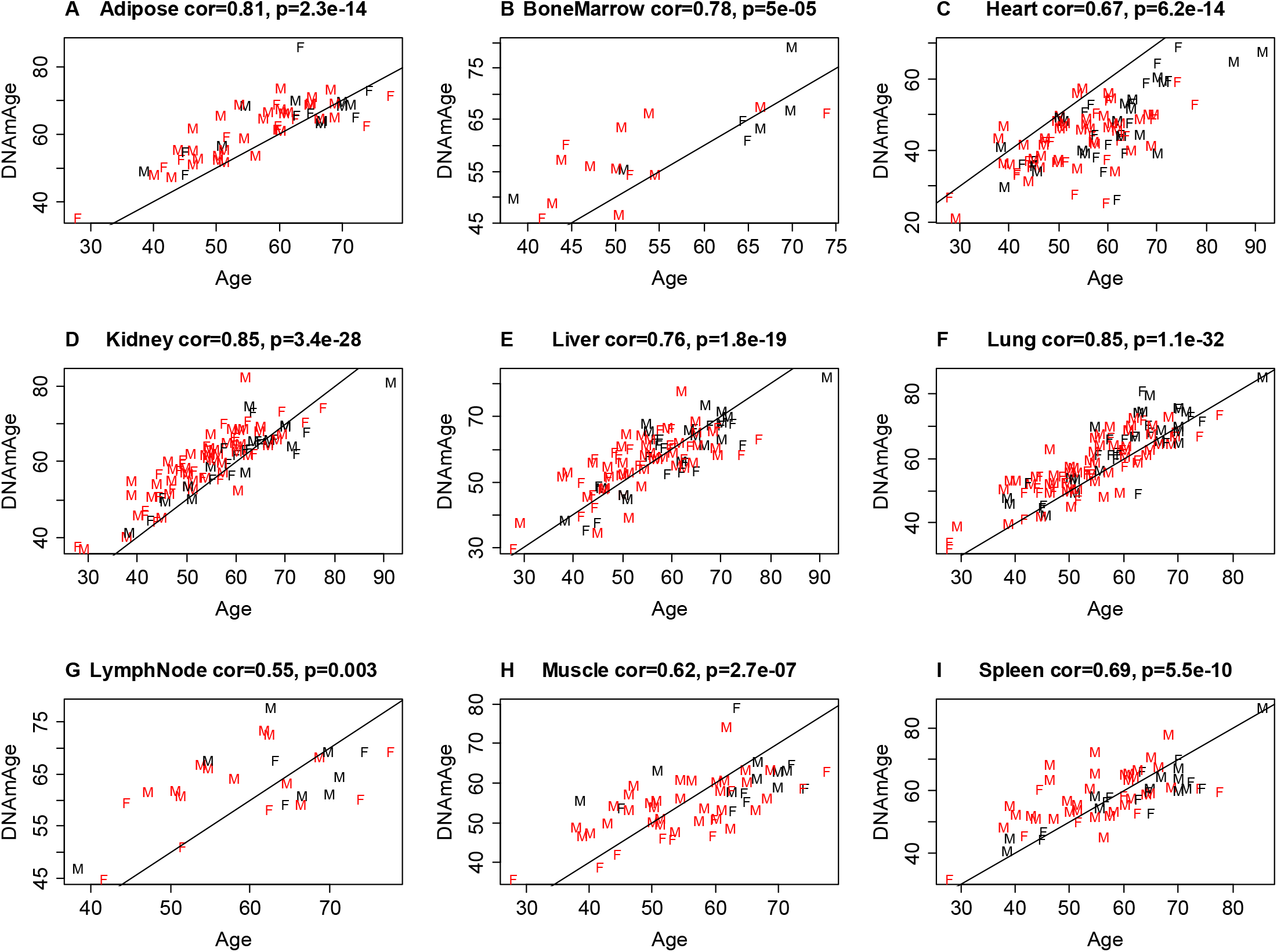
Pan tissue clock applied to different tissues where points are colored by HIV status. This figure is similar to Figure 1 but dots are colored by HIV status. The blood data were omitted since they involved only a single HIV positive individual.

